# The restoration species pool for restoring tropical landscapes: assessment of the largest Brazilian supply chain

**DOI:** 10.1101/568873

**Authors:** Cristina Yuri Vidal, Rafaela Pereira Naves, Ricardo Augusto Gorne Viani, Ricardo Ribeiro Rodrigues

## Abstract

Brazil has been committed to fulfill international restoration goals and to enforce environmental legislation that will require private landowners to undertake ecological restoration of 21 million hectares of degraded and deforested landscapes. To support a broad range of restoration practices, a consolidated supply chain able to represent regional plant diversity is essential. This study investigated the restoration species pool in native plant nurseries in São Paulo state, southeastern Brazil and evaluated their geographic distribution, similarity of their plant stocks and the proportion of species represented from regional floras. Despite the lack of technical assistance and the large presence of non-native species (126 species, average 7.5 species/nursery), we found still more impressive native species richness in plant nurseries (561 species, average 86.4 species/nursery) from both the Atlantic Forest and Cerrado domains, representing 38 to 44% of regional floras. There was a huge bias toward tree and shrub species (96.6%) and absence or underrepresentation of other growth forms, as well as of savanna specialists, animal-dispersed and threatened species. The great dissimilarity of species offered in the nurseries surveyed underscores the importance of regional seed collection practices. Effective assistance and capacitation are essential to address issues related to misidentification of species, underrepresentation of most functional plant groups, and the presence of non-native species, as well as to support the supply chain, currently undergoing market downturn.

**Author contributions:** ‘CYV and RRR conceived and designed the research. CYV performed data compilation; CYV and RPN analyzed the data; CYV and RAGV led the writing of the manuscript; all authors contributed to the drafts and gave final approval for publication.

**Implications for practice:**

- Plant nurseries collecting propagules from the surrounding vegetation provide an adequate – but limited – restoration species pool, with very dissimilar plant stocks available among plant nurseries.
- Plant nurseries concentrate their production on shrub and tree species and sub-represent other growth forms and some functional groups such as animal-dispersed and threatened species.
- The diversity of the restoration species pool is the basis to support a broad range of restoration practice, being essential to boost restoration initiatives that complement and support the conservation of remaining diversity in human modified landscapes.
- Effective assistance and capacity building should be provided to address issues related to misidentification, underrepresentation of functional groups and the presence of exotic and invasive species, as well as to support the supply chain, currently under market downturn

## INTRODUCTION

Recent studies argue it is unlikely that tropical countries will be able to achieve their international commitments to restore ecosystems without spontaneous and assisted regeneration (Chazdon & Uriarte 2016; Crouzeilles et al. 2017), which are less costly than restoration plantings and therefore crucial to scale-up restoration efforts (Holl & Aide 2011; Melo et al. 2013; Brancalion et al. 2016b; Latawiec et al. 2016). However, in landscapes with a long history of land conversion, deforestation and defaunation, resilience is low (Rodrigues et al. 2009; Brancalion et al. 2012a; Bello et al. 2015; Crouzeilles et al. 2017) and vegetation recovery depends on active restoration through direct seeding or planting seedlings (Aerts & Honnay 2011; Holl & Aide 2011; Crouzeilles et al. 2017; Holl 2017; Meli et al. 2017). Indeed, planting trees is the most common tropical forest restoration technique, despite being expensive, time consuming, and labor-intensive (Rodrigues et al. 2011; Palma & Laurance 2015; Brancalion et al. 2016b; Holl et al. 2017; Meli et al. 2017; Jalonen et al. 2018).

Restoration of tropical forests within severely deforested scenarios is a major challenge because seed hand-collection and seedling production is a bottleneck, particularly when intending to represent a large pool of native species and genotypes (Brancalion et al. 2012a; Nevill et al. 2016; Jalonen et al. 2018). There are 40 to 53 thousand tree species within the tropics (Slik et al. 2015) and at least 30 thousand seed plant species in Brazil (Forzza et al. 2012; BFG 2015); as expected, there is not enough knowledge on their biology and current distribution. The challenge goes further when considering the process of harvesting propagules for viable seeds and seedlings’ production, restrained by the reduced and degraded forest cover, lack of information on species reproductive biology and phenological patterns, unskilled labor and deficient technical capacity and assistance (Gregorio et al. 2004; Viani & Rodrigues 2009; Brancalion et al 2012a; Palma & Laurance 2015; Dedefo et al. 2017; Jalonen et al. 2018; White et al. 2018). Despite these setbacks, representation of regional plant diversity is essential to consolidate a native plant market offering an adequate restoration species pool (i.e. native species available in plant nurseries for restoration purposes) (Ladoucer et al. 2017), enabling a broad range of restoration goals (Brancalion et al. 2012b).

Brazil has set a role model regarding restoration initiatives (Aronson et al. 2011; Calmon et al. 2011; Brancalion et al. 2013; Melo et al. 2013; Chaves et al. 2015; Holl 2017; Viani et al. 2017). The country has been committed to fulfill international restoration goals and to enforce a recently revised environmental legislation (i.e., Native Vegetation Protection Law no. 12.651/2012, hereafter NVPL) that applies on private lands, and defines the proportion of native vegetation that must be maintained under protection or restricted use (Brancalion et al. 2016a). To comply with NVPL in case of vegetation deficit, landowners are required to restore vegetation using native species under an ecological restoration perspective (SER 2004). In a few specific cases, landowners are allowed to combine native and non-native species and generate income from the exploitation of their economic potential for timber and non-timber products (Brancalion et al 2012b and 2016a; Amazonas et al. 2018; Cerullo & Edwards 2018). Estimates on NVPL’s restoration demand reaches 21 million hectares (Soares-Filho et al. 2014). Considering that landowners do not collect and produce their own seedlings for active restoration, building up capacity and a supply chain to meet this demand is a major challenge, common to many other tropical countries worldwide.

In fact, most examples of established supply chain for native species aiming large-scale restoration are for non-tropical ecosystems (Ladouceur et al. 2017; Jalonen et al. 2018; White et al. 2018). As an exceptional example, São Paulo state, Brazil developed a supply chain to fulfill their ecological restoration goals, with notable advances on the establishment of plant nurseries during the last 30 years (Barbosa et al. 2003; Martins 2011; Silva et al. 2015, 2017), following the implementation of a legal framework for ecological restoration (Durigan et al. 2010; Aronson et al. 2011; Chaves et al. 2015). Besides, São Paulo state represents an unique opportunity for a case study because i) it is composed by two of the hottest global hotspots, Atlantic Forest and Cerrado (Myers et al. 2000; Forzza et al. 2012) and ii) about 75% of the state has vegetation cover below the 30% threshold (Pardini et al. 2010; Estavillo et al. 2013; Banks-Leite et al. 2014), reinforcing the demand on active restoration. Even though previous assessments and reports investigated plant nurseries’ structure, production capacity and related difficulties (Barbosa & Martins 2003; Martins 2011; Silva et al. 2015, 2017), little or none attention has focused on the composition of the restoration species pool and its ecological aspects regarding taxonomic and functional approaches.

This study evaluated the restoration species pool available in native plant nurseries in São Paulo State, Brazil. We investigated their production capacity regarding number of seedlings, with information about richness and abundances distributions among available species. We also examined the geographical distribution of native plant nurseries along the state, the compositional similarity of their plant stocks and the representation of restoration species pool compared to regional floras. Finally, we explored the relation of diversity descriptors with possible explanatory variables such as production capacity, surrounding forest cover and number of vegetation types. We predicted native species composition is similar among plant nurseries and that they have an average richness around 80 species, in accordance to state recommendations established since 2008 (see details in Aronson et al. 2011; Chaves et al. 2015). We also predicted a compatible but limited representation of regional floras – both under taxonomic and functional approaches - and a positive influence of production capacity, surrounding forest cover and number of vegetation types on overall nurseries’ diversity.

## METHODS

### Data surveys and sampling

We compiled all São Paulo state plant nurseries listed on previous official assessments (Barbosa & Martins 2003; Martins 2011; Silva et al. 2015) plus new or unlisted nurseries indicated by restoration practitioners, totaling 347 plant nurseries. While contacting them by email/website, telephone or in person, we discharged duplicates (n=18), those we could not reach by any means after five attempts (n=55), or that do not produce native species (n=33). Considering 241 eligible plant nurseries, we divided our sample in: i) quick surveys to assess their current production (2015 to 2017); ii) detailed surveys to assess relevant information on the origin of propagules, infrastructure, and market related issues (questionnaire adapted from Oliveira & Zakia 2010), as well as their species and abundance production (raining season 2015/2016) (Appendix S1, Supporting Information).

For the quick surveys, we considered all regions of the São Paulo state, while for detailed surveys we sampled regions with mean forest cover below 30% (i.e. Southwest, Northwest, Center, Southeast) (Fig. 1), where active restoration is usually recommended (Holl & Aide 2011; Tambosi et al. 2013). Additionally, detailed surveys’ sampling depended on plant nurseries’ willingness to provide the requested information.

**Figure 1.**
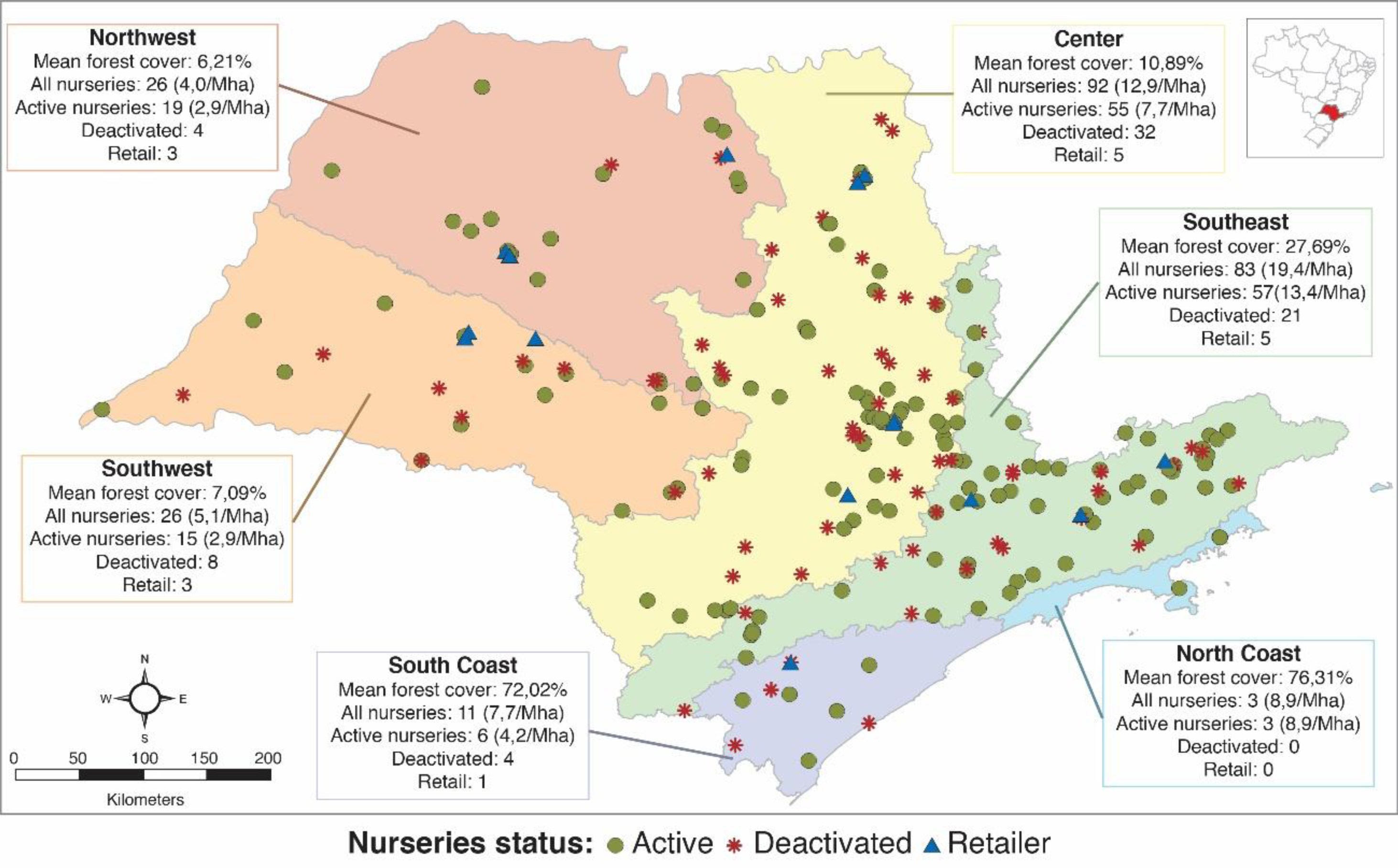
Distribution of native plant nurseries among the ecological regions defined by São Paulo state’s Botanical Institute (SP-IBt). Each region is described 2 by their mean forest cover and by the quantity and density of all assessed native plant nurseries and active nurseries only, as well as the quantity of deactivated 3 and retailing nurseries. Mean forest cover based on São Paulo State Forest Inventory (2011) and density calculated by million hectares (Mha).

Data on species abundance included nursery-grown seedlings available for planting in the field, regardless of the plant container or size. We disregarded hand-collected seeds because plant nurseries usually do not sell them for restoration purposes, but rather use them to produce seedlings. We emphasized the list of available species should consider only those appropriate for ecological restoration projects (i.e. excluding urban afforestation, silviculture, etc.), giving the nursery’s staff free will to choose native species based on their judgment - a common real life practice that can mistakenly lead to the misuse of non-native species.

We dismissed morphospecies identified only to the family or genus levels and standardized species names using the Plantminer tool (www.plantminer.com, Carvalho et al. 2010), according to Flora do Brasil 2020 (http://reflora.jbrj.gov.br/) and The Plant List (www.theplantlist.org). From Flora do Brasil 2020 we retrieved information on growth forms, occurrence (Atlantic Forest and/or Cerrado) and origin (native, non-native), with further evaluation of problematic non-native species according to Sartorelli et al. (2018). For species occurring in the Cerrado (Brazilian Savanna), we refined the classification of occurrence according to their habitat preferences: i) savanna specialist, ii) forest specialist and iii) generalists (Mendonça et al. 2008; Abreu et al. 2017). For a functional grouping approach, we classified native species into the following functional guilds: i) pioneer, ii) fast-growing shading (Rodrigues et al. 2009), iii) understory non-pioneer and iv) canopy non-pioneer. Additionally, species were classified by dispersal syndromes and sub-syndromes (Bello et al. 2017).

For each plant nursery, we calculated the percentage of forest cover and the number of vegetation types (ordinal) in a surrounding 100 km buffer, extracting the information available on official shapefiles provided by the São Paulo State Forest Inventory (2011) with ArcGIS software (University of Campinas license). The six vegetation types occur both within Atlantic Forests (Seasonal Semideciduous Forests (SSF), Atlantic Forest *sensu stricto* (AFSS), Mixed Temperate Araucaria Forests (MTAF), Alluvial and Swamp Forests (A/SF)) and within Cerrado (Cerrado *sensu stricto* and Cerradão).

### Data analysis

We compared native species and families available on plant nurseries with two references. The first reference is the list of species officially recommended for restoration in different regions of São Paulo State, provided by the state’s Botanical Institute (hereafter SP-IBt) and available at www.botanica.sp.gov. The second reference is a dataset of floristic surveys performed by the Forest Ecology and Restoration Laboratory (LERF/University of São Paulo), describing the occurrence of shrub/tree species across forest fragments (N=371) in the studied regions (Rodrigues et al. 2011). Comparisons focused on evaluation of shared and exclusive species, proportions of functional guilds and ranking the botanical families’ richness, in order to detect eventual mismatches or lacking groups in plant nurseries.

To describe the diversity among plant nurseries and within ecological regions we used species abundance distribution models (SAD) (McGill et al. 2007). SAD models provide a powerful way to understand the abundance structure of nurseries’ production, revealing the evenness (Magurran 2013) of their plant stocks. We fitted log series and Poisson log normal distributions to the species abundance data using the maximum-likelihood tools with the sads package for R 3.1 version (Prado et al. 2016). We compared the models based on Akaike’s information criterion (AIC) (Hilborn & Mangel 1997) and for every plant nursery, Poisson log normal provided the best fit to our data. Therefore, we used its parameter sigma (σ) as a local diversity metric (Sæther et al. 2013).

We fitted linear models to analyze the influence of explanatory variables - production capacity, forest cover, number of vegetation types, ecological regions - on richness and sigma diversity descriptors. We tested all models to meet assumptions of normality and homogeneity of variance and then compared them based on AIC. We ranked the models according to the lowest AIC value; models with a difference in AIC (Δ) ≤ 2 can be considered to have equivalently strong empirical support and similar plausibility (Hilborn & Mangel 1997; Bolker 2008).

To evaluate spatial variation of plant stocks’ diversity among studied nurseries, we calculated a multi-site Sorensen dissimilarity index (β_SOR_) as a measure of total β diversity for each region of the state. Then, we calculated the contribution of the turnover and nestedness components of total β diversity, where turnover indicates the dissimilarity resulting from species replacement among plant stocks, whereas nestedness indicates the dissimilarity resulting from differences on species richness (Baselga 2010; Socolar et al. 2016). For practical purposes, high turnover means that plant nurseries have very dissimilar plant stocks in terms of species composition, indicating that regional restoration species pool result from their plant stocks combined. On the other hand, highly nested β diversity means that species-poor plant nurseries have species that are included in the species-rich plant nurseries. For total β diversity (β_SOR_) decomposition, we calculated the Simpson index (β_SIM_) that only measures turnover and the nestedness (β_NEST_) component as the difference of β_SOR_ - β_SIM_ (Baselga 2010). We performed total β diversity (β_SOR_) decomposition using the function “beta-multi.R” in the R package “betapart” (Baselga & Orme 2012).

## RESULTS

### Plant nurseries assessment

We contacted 241 registered native plant nurseries with a “quick” survey, and confirmed that 64.3% (n=155) were still active, while 35.7% were either currently deactivated (n=69) or retailing seedlings from other nurseries (n=17). Geographic distribution of active nurseries was concentrated in the Center (n=55) and Southeast (n=57) regions of São Paulo state (Fig. 1); combined they constituted 72.3% of all plant nurseries.

From detailed surveys made with 54 plant nurseries, we noted that 87% (n=47) collected their own propagules (i.e., seeds and/or fruits), harvesting from the surrounding forest fragments within an average distance of 100 km radius. Over half of them (57%, n=31) also purchased additional seeds from other sources - even from out of the state - to enhance diversity. In the rainy season of 2015/2016 the nurseries we surveyed produced approximately 9.3 million seedlings, with individual production ranging from 3,500 to 1,800,000 seedlings. Regarding identification practices, most active nurseries (90%) kept track of species using both popular and scientific names. However, for botanical identification they relied on their own expertise and/or illustrated guides such as “*Brazilian Trees*” (Lorenzi 2002), as most of them lack access to qualified botanical experts.

### Species diversity

From 687 plant species identified in the nursery survey, 561 (81.1%) were native to São Paulo and 126 (17.8%) were non-native (Tables S1 & S2, Supporting Information). Among natives, there were 542 shrub/tree species (96.6%), five sub-shrubs, seven palm trees and seven lianas, with average richness of 86.4 per nursery, ranging from 18 to 194 species (Table 1). The 126 non-native species represented 4.8% of the total number of seedlings, with particular concern over 10 non-native species that should be avoided in restoration projects due to their invasive potential (Sartorelli et al. 2018) (Table S2, Supporting Information).

**Table 1.** Native and non-native species richness registered for different regions of the of São Paulo state, where we sampled N plant nurseries. Threatened species included extinct, extinct in the wild, critically endangered, endangered or vulnerable species according to IUCN or the State of Sao Paulo’s red list. Regions: SE= southeast, CT=center, NW=northwest, SW=southwest.

| Regions | N | Native species |  |  |  |  | Non- native species |  |  |  |
| --- | --- | --- | --- | --- | --- | --- | --- | --- | --- | --- |
|  |  | all nurseries | min | mean | max | threatened species | all nurseries | min | mean | max |
| SE | 17 | 326 | 18 | 74.4 | 124 | 12 | 44 | 1 | 5.4 | 18 |
| CT | 27 | 472 | 29 | 92.5 | 194 | 16 | 87 | 1 | 8.3 | 22 |
| NW | 6 | 227 | 27 | 87.4 | 116 | 7 | 40 | 2 | 10.2 | 27 |
| SW | 4 | 193 | 54 | 95 | 122 | 7 | 21 | 4 | 7.5 | 11 |
| Total | 54 | 561 | 18 | 86.4 | 194 | 19 | 126 | 1 | 7.5 | 27 |

Considering only native shrub/tree species, plant nurseries’ production encompassed 419 species (37.9%) recommended for restoration by the São Paulo state and 399 species (44.2%) registered within surveys in São Paulo forest remnants, as well as 86 native species that unmatched these floristic lists (Fig. 2A). From all native species available in the plant nurseries, 462 occur in the Atlantic Forest biome and 396 species in the Cerrado, but for the latter, only 94 were savanna specialist species (23.7%), while 250 were forest specialists or generalists (63.1%).

**Figure 2.**
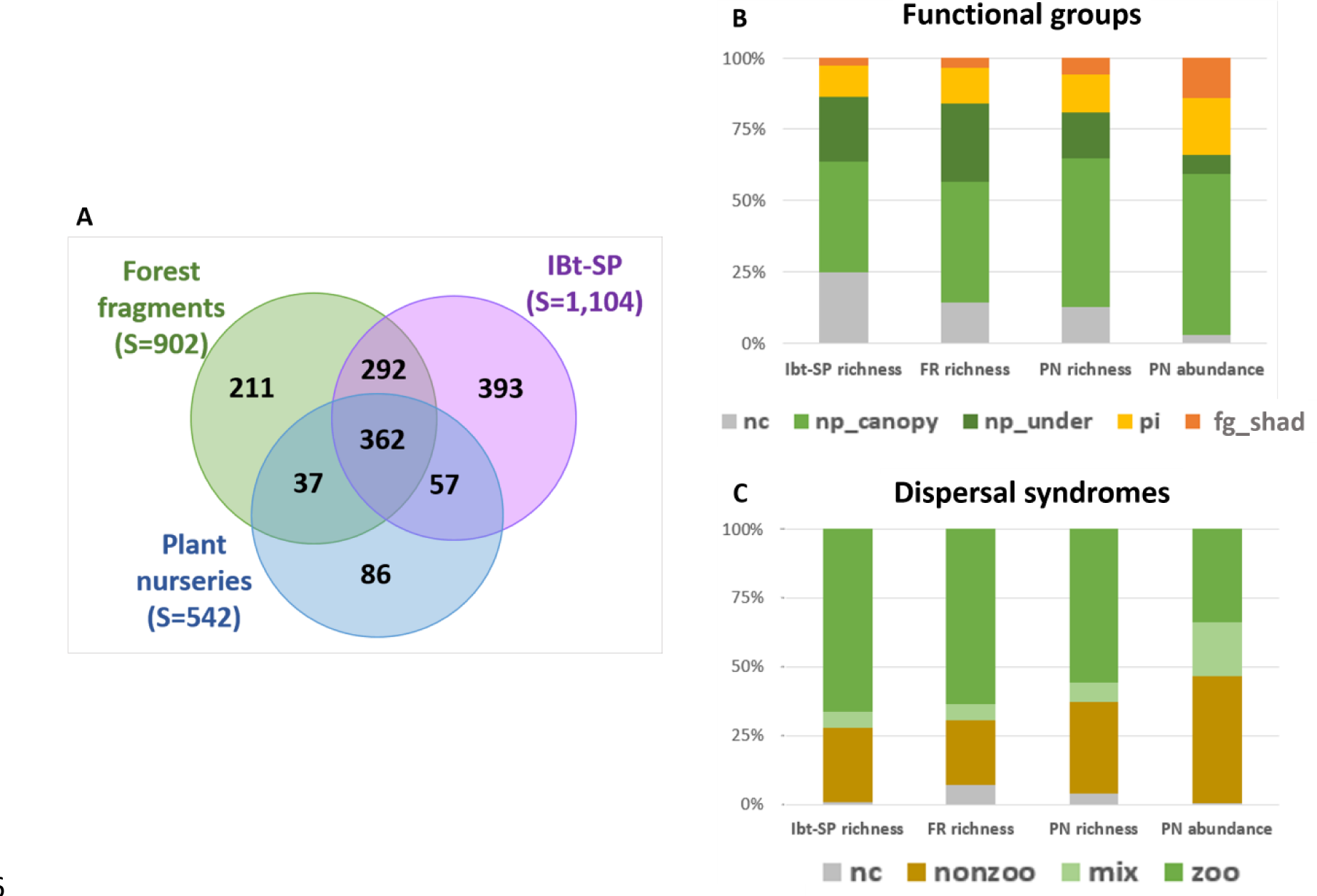
Comparison of floristic composition among plant nurseries (PN), forest fragments (FR) and IBt-SP. (A) Shared and exclusive species richness; (B) Proportion of functional groups/guilds: nc = non-classified, np_canopy = canopy non-pioneer, np_under = understory non-pioneer pi = pioneer and fg_shad = fast-growing shading species; (c) Proportion of dispersal syndromes: nc= non-classified, nonzoo = non zoochoric, mix= mixed (both non-zoo and zoo), zoo = zoochoric species.

Plant nurseries and their plant stocks partially represented species richness and overall proportions of functional groups and dispersal syndromes observed on references (Figs. 2B, 2C). When considering plant stocks’ species abundances, non-pioneer (canopy and understory) were approximately two times more abundant than pioneer and fast-growing shading species (Fig. 2B), while animal-dispersed species represented over a half (56%) of species and one third (34%) of produced seedlings (Figure 3C). Comparison of the richest plant families registered on references with those available on plant nurseries indicated that despite a fair representation of many families, some of them were under-represented – for instance, Lauraceae, Melastomataceae and Rubiaceae families had, on average, less than 30% of their species available on plant nurseries (Figure S1, Supporting Information).

**Figure 3.**
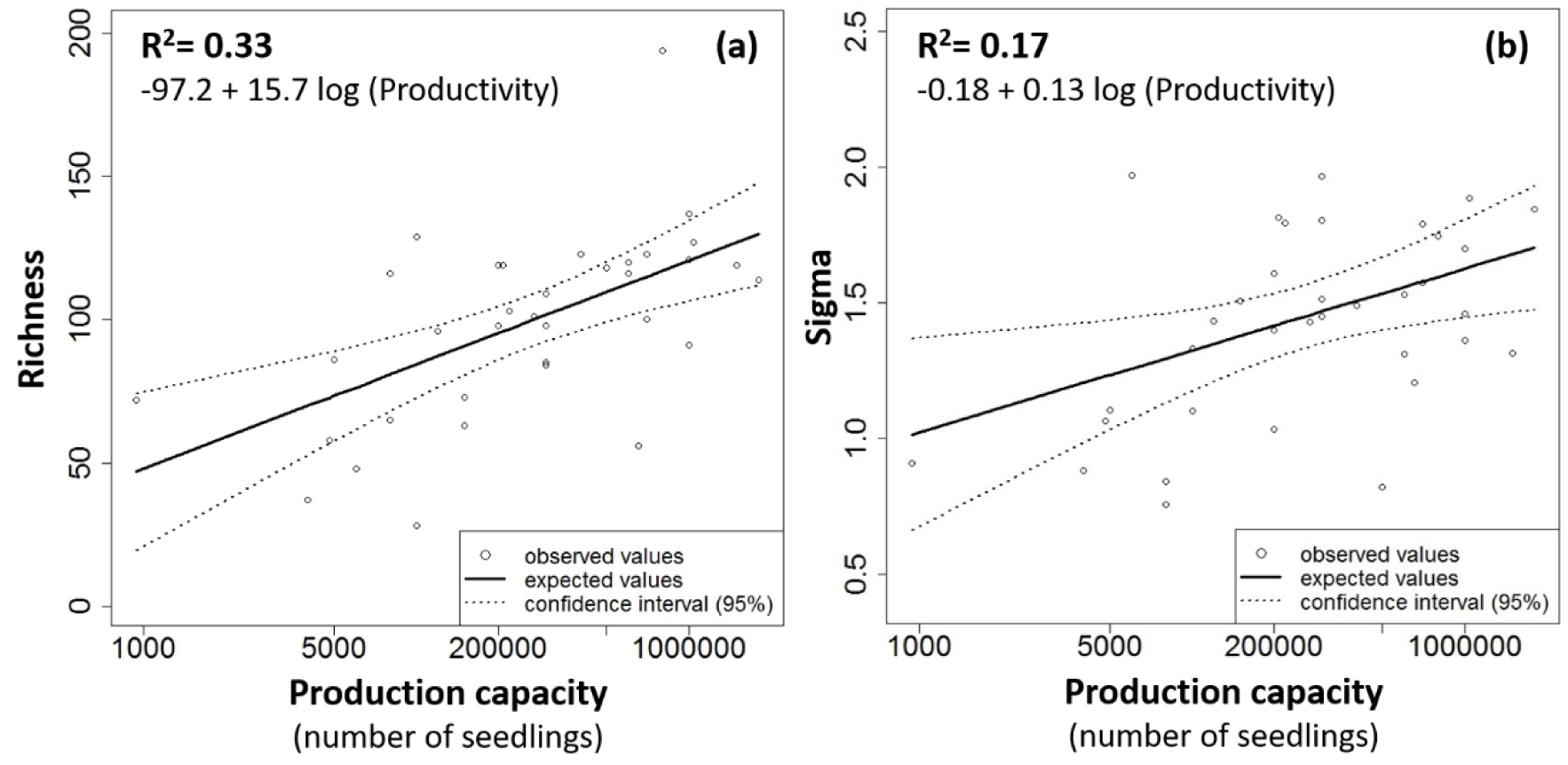
Best fitting linear models for diversity descriptors considering production capacity as an explanatory variable for (a) richness (R^2^ = 0.33) and (b) sigma (R^2^ = 0.34).

Species abundance distribution patterns revealed that nurseries present uneven plant stocks, with 35 species (6.2%) representing half of all seedlings, while the other half included 526 species (93.7%) Regarding species’ frequency among plant nurseries, we classified only 12 species (2%) as common (i.e., occurring in more than 75% nurseries), and 440 species (78.6%) as rare (i.e., occurring in less than 25% nurseries). The 35 most abundant species – 12 of which are also among the most frequent – represent half of seedlings available. They were mostly non-pioneer canopy (23 species), with five fast-growing shading and five pioneer species, and predominantly abiotic-dispersed species (19), with 10 dispersed either by abiotic or biotic factors; only six species are strictly dependent on animals.

In agreement with the above-cited prevalence of rare species, the multi-site Sorensen dissimilarity index (β_SOR_) presented high values in all regions of the state, with a consistent major contribution from its turnover component, revealed by the higher values of β_SIM_ (Table 2).

**Table 2.** Total beta diversity (β_SOR_) decomposed into turnover (β_SIM_) and nestedness (β_NEST_) components for all distinct regions. N is the number of sampled plant nurseries.

| Regions | N | $\beta_{SOR}$ | $\beta_{SIM}$ | $\beta_{NEST}$ |
| --- | --- | --- | --- | --- |
| Southeast | 17 | 0.87 (100%) | 0.78 (89.6%) | 0.09 (10.4%) |
| Center | 27 | 0.90 (100%) | 0.83 (92.2%) | 0.07 (7.8%) |
| Northwest | 6 | 0.70 (100%) | 0.58 (82.8%) | 0.12 (17.2%) |
| Soutwest | 4 | 0.58 (100%) | 0.45 (77.5%) | 0.13 (22.5%) |
| Total | 54 | 0.95 (100%) | 0.91 (95.8%) | 0.04 (4.2%) |

There was a positive correlation between the production capacity of a plant nursery and its species richness and sigma diversity (Fig. 3). Models considering forest cover, number of vegetation types, ecological regions were no better than expected by chance (null model) (Table S3, Supporting Information).

## DISCUSSION

Our study on the largest native supply chain in Brazil (Silva et al. 2017) revealed that the restoration species pool offered approximately 38% of native tree and shrub species recommended for restoration in São Paulo state. We registered high overall native species richness (561) and high average species richness per nursery (86.4), which is above Brazilian national standards (63) (Silva et al. 2015) and above previous estimates for plant nurseries in the state (Barbosa et al. 2003; Martins 2011). Another remarkable result of our study is the singularity of plant nurseries’ production, proven by the high values of β-diversity due to its turnover component (i.e., high dissimilarity among nurseries’ plant stocks). Since most plant nurseries collect propagules from the surroundings, we presume that not only they represent the regional taxonomic diversity but also the populations’ genotypes (Zucchi et al. 2017; White et al. 2018). These results altogether reinforce the importance of the restoration supply chain, especially in those regions where restoration cannot rely on spontaneous regeneration processes and where a well-established native plant market may contribute to high diversity ecological restoration initiatives.

Despite the remarkable diversity of the restoration species pool we studied, the representation of regional floras was biased under several aspects. While both Atlantic Forest and Cerrado species were available in plant nurseries, there was an underrepresentation of savannas’ specialists, which may lead to afforestation and to other negative consequences for the native diversity of the grassy biomes of Cerrado (Overbeck et al. 2013; Veldman et al. 2015a, 2015b; Abreu et al. 2017; Buisson et al. 2018). Furthermore, the restoration species pool constituted a narrow spectrum of growth forms that lack or underrepresent lianas, epiphytes and herbs; in the Atlantic Forest and Cerrado biomes, these growth forms exceed 2 to 7 times the number of tree species (BFG 2015). Although production bias towards woody species exists because they are the main structural components of forests – the main target of Brazilian restoration initiatives - awareness should be raised as to the importance of other growth forms, especially for non-forest biomes, where restoration demand is increasing and propagation knowledge is still challenging (Overbeck et al. 2013; Campbell et al. 2015; Veldman et al. 2015a; Garcia et al. 2016; Mayfield 2016; Buisson et al. 2018). Considering only tree and shrub species, plant nurseries were offering customers less than 50% of species registered on studied references, reinforcing it is a huge challenge to offer species diversity for tropical diverse ecological restoration, even in the most established supply chain in Brazil (Silva et al. 2015, 2017). The situation is even more critical when considering that only 2.3% (19 species) of São Paulo state’s threatened plant species were found in plant nurseries, falling short of the objectives of the Global Strategy for Plant Conservation in Brazil, which defines a goal of making 20% of threatened species available for restoration efforts by 2020 (Martins et al. 2017). Since threatened species offer a greater challenge for conservationists, specific recovery plans would be necessary to achieve this particular goal (Durigan et al. 2010; Martins et al. 2017).

A positive aspect we highlight is that among the functional groups available in plant stocks, there were fast-growing shading tree species that boost soil coverage and shade exotic weeds (Rodrigues et al. 2009), as well as a great variety and quantity of canopy non-pioneer species, which will presumably persist in restored sites over time (Rodrigues et al. 2011; Brancalion et al. 2012a). However, the overall variety and quantities of animal-dispersed species are below those expected for tropical forests, which varies from 70 to 94% of woody species (Almeida-Neto et al. 2008; Bello et al. 2017). As shown by Brancalion et al. (2018), large-seeded animal-dispersed species are particularly underrepresented in restoration projects, with consequences for carbon storage and restoration outcomes. We recommend enhancement on the proportion of animal-dispersed species in plant nurseries, since plants consumed and dispersed by animals are notably important in degraded landscapes, where maintenance of plant-animal interactions are essential to enable restoration of ecological processes, biological fluxes and ecosystem services (Howe 2016; Brancalion et al. 2018).

Tropical forests are typically characterized by skewed species-abundance distributions, with few common species and many rare or very rare (Caiafa & Martins 2010; Hubbell 2013). As a possible reflection of this pattern, we found that almost 80% of species were rare among plant nurseries, and half of total seedling production was composed by only 35 out of 561 native species (6%). These results are consonant with our findings on the high dissimilarity among plant stocks (i.e., high total B diversity), mainly due to the replacement of species among them (i.e., turnover) (Baselga 2010; Socolar et al. 2016). Previous studies have consistently indicated turnover as the larger component of total β-diversity in tropical ecosystems (Soininen et al. 2018), a pattern that was registered within the Atlantic Forest Domain in São Paulo state (Bergamin et al. 2017, Farah et al. 2017). Therefore, one possible explanation for the high dissimilarities among plant stocks may be related to plant nurseries’ practice of collecting propagules from surrounding forest fragments, which are described as highly variable regarding species composition (Bergamin et al. 2017, Farah et al. 2017). In this sense, well-distributed nurseries not only maximize the chances of representing local specimens adapted for regional restoration projects (White et al. 2018) but may also enhance the taxonomic representation of regional floras. Thus, the biased geographic distribution of plant nurseries raises an issue to be addressed by public policy makers: a better regional planning must align restoration demand and seeds and seedlings production, and foster corrective and supportive measures such as the implementation of inter-regional seed exchange programs (Brancalion et al. 2012a; Jalonen et al. 2018).

Although plant stocks were assembled from surrounding fragments, we did not find evidence supporting the influence of the percentage of surrounding forest cover and number of vegetation types over the restoration species pool’s diversity. That is probably because all nurseries evaluated on the detailed surveys were located in regions with reduced forest cover (less than 30%) and with little variation on vegetation types. Beyond the positive influence of production capacity over the diversity of the restoration species pool, we consider that overall, high species richness most likely derived from the enforcement of São Paulo state legislation (Brancalion et al. 2010). Until 2014, the state legislation used to establish that restoration projects should reach a minimum of 80 native woody plant species (Aronson et al. 2011); currently, it has shifted the focus from the number of reintroduced species to the monitoring of structure and diversity goal achievement (Chaves et al. 2015). Regardless of the discussion on whether it is positive or negative to standardize the amount of species on a restoration project (Aronson et al. 2011), we must recognize that these legal instruments have pushed plant nurseries to enhance their diversity (Brancalion et al. 2012a; Silva et al. 2017), placing São Paulo state native trees’ seedling production at a very high level, far higher than elsewhere in Brazil, and possibly worldwide.

Despite the positive aspects we detected on the diversity of the restoration species pool, we must consider some caveats. First, we highlight the worrisome market downturn that have been affecting the production of native seedlings since the initial discussions to revise the main environmental legislation in Brazil (i.e., Native Vegetation Protection Law no. 12.651/2012) (details in Brancalion et al. 2016a). Second, we considered only the rainy season of 2015/2016 and species richness may be even higher if a longer period is evaluated, as flowering and fruiting periods have interannual variability (Morellato et al. 2001; Viani & Rodrigues 2009; Brancalion et al. 2012a). Third, few nurseries adopt good identification practices such as the collection of voucher specimens for depositing in herbaria and examination by professional botanists. Mistaken identification in plant nurseries can mislead to over- or under-estimations of the actual diversity available on nurseries and it may also explains the production of non-native species, a common issue in restoration sites (Barbosa et al. 2003; Assis et al. 2013; Brancalion et al. 2016b).

Our results underscore that the restoration species pool in São Paulo comprehends a considerable portion of tree and shrub diversity, but how it affects success of ecological restoration depends on whether we consider biodiversity introduced in restoration projects as a goal or a driver of the recovery process (Naeem 2016). There is an ongoing debate in Brazil regarding the benefits of using high or low diversity in restoration efforts (Brancalion et al. 2010; Durigan et al. 2010; Aronson et al. 2011). It considers arguments related to cost reductions, field performance and the definition of presumed “framework species” (Suganuma & Durigan 2015), as well as compelling evidence associating biodiversity and ecosystem functioning (BEF) (Aerts & Honnay 2011; Cardinale et al. 2012; Tilman et al. 2014; Brockerhoff et al. 2017). Despite the lack of consistent evidence relating the amount of reintroduced diversity and restoration success, pursuing and promoting higher diversity in the restoration species pool is essential to broaden and foster a wide variety of restoration approaches. A large restoration species pool could benefit other conservation purposes such as the restoration of degraded forest remnants (Viani et al. 2015), and enable some economic trade-off for landowners who comply with the law, through the sustainable exploitation of timber and non-timber products from native species (Brancalion et al 2012b and 2016a; Amazonas et al. 2018; Cerullo & Edwards 2018).

The impressive levels of species richness registered in this study represent, to our knowledge, the most diverse tropical native tree seedling production and supply chain anywhere in the world. Particularly on human-modified landscapes with reduced forest cover, plant nurseries play a pivotal role propagating the remaining biodiversity, as they collect most of their seed from local provenances and represent local populations and communities (Jalonen et al. 2018). However, even a well-established supply chain offered a restrained restoration species pool, limited by deficient knowledge on species’ biology, uneven plant nurseries’ geographic distribution, and lack of technical capacitation and assistance. These limitations expose issues and opportunities to be addressed by restoration policies aiming to optimize the full potential of restoration plantings, especially supporting the conservation value of forest fragments in human-modified landscapes.

## ACKNOWLEDGEMENTS

We thank all plant nurseries’ owners and staff for sharing their information and thoughts; Letícia S. Santos, Bruno H. Guastala, Fernando H. Silva and Sergio Lozano-Baez for contacting nurseries during quick surveys. We warmly thank James Aronson for comments and suggestions that helped to improve this manuscript. This study was financed in part by the Coordenação de Aperfeiçoamento de Pessoal de Nível Superior – Brasil (CAPES) – Finance Code 001, by the National Council for Scientific and Technological Development (CNPq grant #870360/1997-3) and by The São Paulo Research Foundation (FAPESP grant # 2013/50718-5).

## LITERATURE CITED

Abreu RCR, Hoffmann WA, Vasconcelos HL, Pilon NA, Rossatto DR, Durigan G (2017) The biodiversity cost of carbon sequestration in tropical savanna. Science Advances 3:e1701284

Aerts R, Honnay O (2011) Forest restoration, biodiversity and ecosystem functioning. BMC Ecology 24:11–29

Aronson J, Brancalion PHS, Durigan G, Rodrigues RR, Engel VL, Tabarelli M, et al. (2011) What Role Should Government Regulation Play in Ecological Restoration? Ongoing Debate in São Paulo State, Brazil. Restoration Ecology 19:690–695

Assis GB, Suganuma MS, Melo ACG, Durigan G (2013) Uso de espécies nativas e exóticas na restauração de matas ciliares no Estado de São Paulo (1957 - 2008). Revista Árvore 37:599–609

Banks-Leite C, Pardini R, Tambosi L, Pearse WD, Bueno AA, Bruscagin RT et al. (2014) Using ecological thresholds to evaluate the costs and benefits of set-asides in a biodiversity hotspot. Science 345:1041–1045

Barbosa LM, Barbosa JM, Barbosa KC, Potomati A, Martins SE, Asperti LM et al. (2003) Recuperação Florestal Com Espécies Nativas No Estado De São Paulo: Pesquisas Apontam Mudanças Necessárias. Florestar estatístico 6:28–34

Barbosa LM and Martins SE (2003) Diversificando o reflorestamento no estado de São Paulo: espécies disponíveis por região e ecossistema. Secretaria de Meio Ambiente do Estado de São Paulo, São Paulo

Baselga A (2010) Partitioning the turnover and nestedness components of beta diversity. Global Ecology and Biogeography 19:134–143

Baselga A, Orme CDL (2012) Betapart: An R package for the study of beta diversity. Methods in Ecology and Evolution 3:808–812

Bello C, Galetti M, Montan D, Pizo MA, Mariguela TC, Culot L, et al. (2017) Atlantic frugivory: A plant – frugivore interaction data set for the Atlantic Forest. Ecology 0:0

Bello C, Galetti M, Pizo MA, Magnago LFS, Rocha MF, Lima RAF, Peres CA, Ovaskainen O, Jordano P (2015) Defaunation affects carbon storage in tropical forests. Science Advances 1:1–11

Bergamin RS, Bastazini VAG, Vélez-Martin E, Debastiani V, Zanini KJ, Loyola R, Müller SC (2017) Linking beta diversity patterns to protected areas: lessons from the Brazilian Atlantic Rainforest. Biodiversity and Conservation

BFG The Brazil Flora Group (2015) Growing knowledge: an overview of Seed Plant diversity in Brazil. Rodriguesia 66:1–29

Bolker BM (2008) Ecological models and data in R. Princeton University Press, New Jersey

Brancalion PHS, Bello C, Chazdon RL, Galetti M, Jordano P, Lima RAF, et al. (2018) Maximizing biodiversity conservation and carbon stocking in restored tropical forests. Conservation Letters:e12454

Brancalion PHS, Garcia LC, Loyola R, Rodrigues RR, Pillar VD, Lewinsohn TM (2016a) A critical analysis of the Native Vegetation Protection Law of Brazil (2012): Updates and ongoing initiatives. Natureza e Conservacao 14:1–15

Brancalion PHS, Melo FPL, Tabarelli M, Ricardo R (2013) Biodiversity persistence in highly human-modified tropical landscapes depends on ecological restoration. Tropical Conservation Science 6:705–710

Brancalion PHS, Rodrigues RR, Gandolfi S, Kageyama PY, Nave AG, Gandara FB, Barbosa LM, Tabarelli M (2010) Instrumentos legais podem contribuir para a restauração de florestas tropicais biodiversas. Revista Árvore 34:455–470

Brancalion PHS, Schweizer D, Gaudare U, Mangueira JR, Lamonato F, Farah FT, Nave AG, Rodrigues R (2016b) Balancing economic costs and ecological outcomes of passive and active restoration in agricultural landscapes: the case of Brazil. Biotropica 48:856–867.

Brancalion PHS, Viani RAG, Aronson J, Rodrigues RR, Nave AG (2012a) Improving Planting Stocks for the Brazilian Atlantic Forest Restoration through Community-Based Seed Harvesting Strategies. Restoration Ecology 20:704–711

Brancalion PHS, Viani RAG, Strassburg BBN, Rodrigues RR (2012b) Finding the money for tropical forest restoration. Unasylva 63:41–50

Brockerhoff EG, Barbaro L, Castagneyrol B, Forrester DI, Gardiner B, Lyver POB et al. (2017) Forest biodiversity, ecosystem functioning and the provision of ecosystem services. Biodiversity and Conservation 26:3005

Buisson E, Le Stradic S, Silveira FAO, Durigan G, Overbeck GE, Fidelis A et al. (2018) Resilience and restoration of tropical and subtropical grasslands, savannas, and grassy woodlands. Biological Reviews

Caiafa AN and Martins FR (2010) Forms of rarity of tree species in the southern Brazilian Atlantic rainforest. Biodiversity and Conservation 19:2597–2618

Calmon M, Brancalion PHS, Paese A, Aronson J, Castro P, Silva SC, Rodrigues RR (2011) Emerging Threats and Opportunities for Large-Scale Ecological Restoration in the Atlantic Forest of Brazil. Restoration Ecology 19:154–158

Cardinale BJ, Duffy E, Gonzalez A, Hooper DU, Perrings C, Venail P, et al. (2012) Biodiversity loss and its impact on humanity. Nature 489:326–326

Carvalho GH, Cianciaruso MV, Batalha MA (2010) Plantminer: A web tool for checking and gathering plant species taxonomic information. Environmental Modelling and Software 25:815–816

Cerullo GR & Edwards DP (2018) Actively restoring resilience in selectively logged tropical forests. Journal of Applied Ecology, 56, 1:107–118

Chaves RB, Durigan G, Brancalion PHS, Aronson J (2015) On the need of legal frameworks for assessing restoration projects success: new perspectives from São Paulo state (Brazil). Restoration Ecology 23:754–759

Chazdon RL and Uriarte M (2016) Incorporating natural regeneration in forest landscape restoration in tropical regions: synthesis and key research gaps. Biotropica 48:915–924

Crouzeilles R, Curran M, Ferreira MS, Lindenmayer DB, Grelle CE V, Rey Benayas JM (2016) A global meta-analysis on the ecological drivers of forest restoration success. Nature communications 7:1–8

Crouzeilles R, Ferreira MS, Chazdon RL, Lindenmayer DB, Sansevero JBB, Monteiro L, et al. (2017) Ecological restoration success is higher for natural regeneration than for active restoration in tropical forests. Science Advances 3:e1701345

Dedefo K, Derero A, Tesfaye Y, Muriuki J (2017) Tree nursery and seed procurement characteristics influence on seedling quality in Oromia, Ethiopia. Forests Trees and Livelihoods 26:96–110

Durigan G (2012) Estrutura e diversidade de comunidades florestais. Pages 294–319 In:Martins SV (editor)Ecologia de florestas tropicais do Brasil. Universidade Federal de Viçosa, Viçosa, Minas Gerais

Durigan G, Engel VL, Torezan JM, Melo ACG, Marques MCM, Martins SV, et al. (2010) Normas jurídicas para a restauração ecológica: uma barreira a mais a dificultar o êxito das iniciativas? Revista Árvore 34:471–485

Estavillo C, Pardini R, Rocha PLB (2013) Forest loss and the biodiversity threshold: An evaluation considering species habitat requirements and the use of matrix habitats. PLoS ONE 8:1–10

Farah FT, Muylaert RL, Ribeiro MC, Ribeiro JW, Mangueira JRSA, Souza VC, Rodrigues RR (2017) Integrating plant richness in forest patches can rescue overall biodiversity in human-modified landscapes. Forest Ecology and Management 397:78–88

Flora do Brasil 2020 Jardim Botânico do Rio de Janeiro. Available on http://floradobrasil.jbrj.gov.br/ (acessed 20 July 2018)

Forzza RC, Baumgratz JF, Bicudo CEM, Canhos DAL, Carvalho Jr AA, Coelho MAN et al. (2012) New Brazilian Floristic List Highlights Conservation Challenges. BioScience 62:39–45

Garcia LC, Hobbs RJ, Ribeiro DB, Tamashiro JY, Santos FAM, Rodrigues RR (2016) Restoration over time: is it possible to restore trees and non-trees in high-diversity forests? Applied Vegetation Science 19, 655–666

Gregorio N, Herbohn J, Harrison S (2004) Small-scale forestry development in Leyte, Philippines: The central role of nurseries. Small-scale Forest Economics, Management and Policy 3:337–351

Hilborn R and Mangel M (1997) The Ecological Detective Confronting Models with Data. Princeton University Press, New Jersey

Holl KD (2017) Restoring tropical forests from the bottom up. Science 355:455–456

Holl KD and Aide TM (2011) When and where to actively restore ecosystems? Forest Ecology and Management 261:1558–1563

Holl KD, Reid JL, Chaves-Fallas JM, Oviedo-Brenes F, Zahawi RA (2017) Local tropical forest restoration strategies affect tree recruitment more strongly than does landscape forest cover. Journal of Applied Ecology 54:1091–1099

Hubbell SP (2013) Tropical rain forest conservation and the twin challenges of diversity and rarity. Ecology and Evolution 3:3263–3274

Jalonen R, Valette M, Boshier D, Duminil J, Thomas E (2018) Forest and landscape restoration severely constrained by a lack of attention to the quantity and quality of tree seed: Insights from a global survey. Conservation Letters

Ladouceur E, Jim B, Marin M, Vitis M De, Abbandonato H, Iannetta PPM, Bonomi C, Pritchard HW (2017) Native Seed Supply and the Restoration Species Pool. Conservation Letters 11(2): e12381. https://doi.org/10.1111/conl.12381

Latawiec AE, Crouzeilles R, Brancalion PHS, Rodrigues RR, Sansevero JB, Santos JS, et al. (2016) Natural regeneration and biodiversity: a global meta-analysis and implications for spatial planning. Biotropica 48:844–855

Lorenzi H (2002) Árvores Brasileiras: manual de identificação e cultivo de plantas arbóreas do Brasil. Instituto Plantarum, Nova Odessa, São Paulo

Magurran AE (2013) Measuring biological diversity. Blackwell Publishing Company

Martins E, Loyola R, Martinelli G (2017) Challenges and Perspectives for Achieving the Global Strategy for Plant Conservation Targets in Brazil. Annals of the Missouri Botanical Garden 102:347–356

Martins RB (2011) Diagnóstico dos produtores de mudas florestais nativas do Estado de São Paulo. Secretaria de Meio Ambiente do Estado de São Paulo, São Paulo

McGill BJ, Etienne RA, Gray JS, Alonso D, Anderson MJ, Benecha HK, et al. (2007) Species abundance distributions: Moving beyond single prediction theories to integration within an ecological framework. Ecology Letters 10:995–1015

Meli P, Holl KD, Benayas JMR, Jones HP, Jones PC, Montoya D, Mateos DM (2017) A global review of past land use, climate, and active vs. passive restoration effects on forest recovery. PLoS ONE 12:1–17

Melo FPL, Pinto SRR, Brancalion PHS, Castro PS, Rodrigues RR, Aronson J, Tabarelli M (2013) Priority setting for scaling-up tropical forest restoration projects: Early lessons from the Atlantic forest restoration pact. Environmental Science and Policy 33:395– 404

Mendonça RC, Felfili JM, Walter BM, Silva Junior MC, Rezende AV, Filgueiras TS, Nogueira P (2008) Flora vascular do bioma Cerrado. Pages 422–442 In Sano SM, Almeida SP, Ribeiro JF (eds.) Flora vascular do bioma Cerrado: checklist com 12.356 espécies. EMBRAPA Cerrados, Brasília, Distrito Federal

Morellato LPC, Talora DC, Takahasi A, Bencke CC, Romera EC, Zipparro VB (2001) Phenology of Atlantic rain forest trees: A comparative study. Biotropica 32:811–823

Myers N, Mittermeier RA, Mittermeier CG, Fonseca GA, Kent J (2000) Biodiversity hotspots for conservation priorities. Nature 403:853–858

Naeem S (2016) Biodiversity as a Goal and Driver of Restoration. Pages 57–89 In Palmer, MA, Zedler JB, Falk DA (eds.) Foundations of Restoration Ecology. Second edition. Island Press, Washington

Nevill PG, Tomlinson S, Elliott CP, Espeland EK, Dixon KW, Merritt DJ (2016) Seed production areas for the global restoration challenge. Ecology and Evolution 6:7490–7497

Oliveira RE and Zakia MJB (2010) Guia para análise de viveiros de mudas nativas - Checklist para verificação da adequação legal, socioambiental e ecológica de viveiros de mudas florestais nativas. Piracicaba, São Paulo

Overbeck GE, Hermann JM, Andrae BO, Boldrini II, Kiehl K, Kirmer A, et al. (2013) Restoration ecology in Brazil-time to step out of the forest. Natureza a Conservacao 11:92–95

Palma AC and Laurance SGW (2015) A review of the use of direct seeding and seedling plantings in restoration: What do we know and where should we go? Applied Vegetation Science 18:561–568

Pardini R, Bueno ADA, Gardner TA, Prado PI, Metzger JP (2010) Beyond the fragmentation threshold hypothesis: regime shifts in biodiversity across fragmented landscapes. PloS one 5:e13666

Prado PI, Miranda MD, Chalom A (2016) sads: Maximum Likelihood Models for Species Abundance Distributions R package version 0.3.1. Available from https://cran.r-project.org/package=sads

Rodrigues RR, Gandolfi S, Nave AG, Aronson J, Barreto TE, Vidal CY, Brancalion PHS (2011) Large-scale ecological restoration of high-diversity tropical forests in SE Brazil. Forest Ecology and Management 261: 1605–1613

Rodrigues RR, Lima RAF, Gandolfi S, Nave AG (2009) On the restoration of high diversity forests: 30 years of experience in the Brazilian Atlantic Forest. Biological Conservation 142:1242–1251

Sæther BE, Engen S, Grøtan V (2013) Species diversity and community similarity in fluctuating environments: Parametric approaches using species abundance distributions. Journal of Animal Ecology 82:721–738

Sartorelli PAR, Benedito ALD, Campos Filho EM, Sampaio AB, Ana S, Gouvêa APML (2018) Guia de plantas não desejáveis na restauração florestal. Agroicone, São Paulo. Available from http://www.inputbrasil.org/wp-content/uploads/2018/03/guia-plantas-nao-desejaveis.pdf

SER Society for Ecological Restoration International Science & PolicyWorking Group (2004) The SER International Primer on Ecological Restoration. Tucson, Arizona.

Silva APM, Marques HR, Santos TVMN, Teixeira AMC, Luciano MSF, Sambuichi RHR (2015) Diagnóstico da Produção de Mudas Florestais Nativas no Brasil. IPEA, Brasília, Distrito Federal

Silva APM, Schweizer D, Marques HR, Cordeiro AM, Nascente TVM, Sambuichi RHR, et al. (2017)Can current native tree seedling production and infrastructure meet an increasing forest restoration demand in Brazil? Restoration Ecology 25:509–515

Slik JWF et al. (2015) An estimate of the number of tropical tree species. Proceedings of the National Academy of Sciences 112:7472–7477

Soares-Filho B, Rajão R, Macedo M, Carneiro A, Costa W, Coe M, et al. (2014) Cracking Brazil ’ s Forest Code. Science 344:363–364

Socolar JB, Gilroy JJ, Kunin WE, Edwards DP (2016) How Should Beta-Diversity Inform Biodiversity Conservation? Trends in Ecology and Evolution 31:67–80

Soininen J, Heino J, Wang J (2018) A meta-analysis of nestedness and turnover components of beta diversity across organisms and ecosystems. Global Ecology and Biogeography 27:96–109

Suganuma MS and Durigan G (2015) Indicators of restoration success in riparian tropical forests using multiple reference ecosystems. Restoration Ecology 23:238–251

Tambosi LR, Martensen AC, Ribeiro MC, Metzger JP (2013) A framework to optimize biodiversity restoration efforts based on habitat amount and landscape connectivity. Restoration Ecology 22:169–177

R Development Core Team (2011) R: a language and environment for statistical computing. R Foundation for Statistical Computing, Vienna, Austria

Tilman D, Isbell F, Cowles JM (2014) Biodiversity and ecosystem functioning. Annual Review of Ecology, Evolution, and Systematics 45:471–493

Veldman JW, Overbeck GE, Negreiros D, Mahy G, Le Stradic S, Fernandes GW, et al. (2015a) Tyranny of trees in grassy biomes. Science 347:484–485

Veldman JW, Overbeck GE, Negreiros D, Mahy G, Stradic SLE, Fernandes GW, et al. (2015b) Where Tree Planting and Forest Expansion are Bad for Biodiversity and Ecosystem Services. BioScience 65:1011–1018

Viani RAG, Holl KD, Padovezi A, Strassburg BBN, Farah FT, Chaves RB, et al. (2017) Protocol for Monitoring Tropical Forest Restoration: Perspectives From the Atlantic Forest Restoration Pact in Brazil. Tropical Conservation Science 10:1–8

Viani RAG, Mello FNA, Chi IE, Brancalion PHS (2015) A new focus for ecological restoration:Management of degraded forest remnants in fragmented landscapes. GPL news November:5–9

Viani RAG and Rodrigues RR (2009) Potential of the seedling community of a forest fragment for tropical forest restoration. Scientia Agricola 66:772–779

White A, Fant J, Havens K, Skinner M, Krammer A (2018) Restoring species diversity: assessing capacity in the United States native plant industry. Restoration Ecology 26: 605–611

Zucchi MI, Suiji PS, Mori GM, Viana JPG, Grando C, Silvestre EA, et al. (2017) Genetic diversity of reintroduced tree populations in restoration plantations of the Brazilian Atlantic Forest. Restoration Ecology 26:694–701

